# Golgi-localized PAQR4 mediates Anti-apoptotic Ceramidase Activity in Breast Cancer

**DOI:** 10.1101/772665

**Authors:** Line Pedersen, Pouda Panahandeh, Muntequa Ishtiaq Siraji, Stian Knappskog, Per Eystein Lønning, Qingzhang Zhu, Ruth Gordillo, Philipp E. Scherer, Anders Molven, Knut Teigen, Nils Halberg

## Abstract

The metabolic network of sphingolipids plays important roles in cancer biology. Prominent sphingolipids include ceramides and sphingosine-1-phosphate that regulate multiple aspects of growth, apoptosis and cellular signaling. Although many significant enzymatic regulators of the sphingolipid pathway have been described in detail, the list is currently incomplete. Here, we applied a systemic approach to identify and molecularly define progestin and adipoQ receptor family member IV (PAQR4) as a Golgi-localized ceramidase. We find PAQR4 to be ∼5 fold upregulated in breast cancer compared to matched control tissue and that its overexpression correlate with disease-specific survival rates in breast cancer. PAQR4 is a seven transmembrane protein, and depletion of PAQR4 leads to cellular apoptosis through accumulation of ceramides. Our findings establish PAQR4 as Golgi-localized ceramidase required for cellular growth in breast cancer.

## Introduction

Sphingolipids are key regulatory bioactive molecules that play important roles in cancer biology (Ogretmen, 2018). Altered sphingolipid metabolism are linked to several cancer types including liver, colon (Selzner et al., 2001), endometrial (Knapp et al., 2010) and breast cancer (Moro et al., 2018; Nagahashi et al., 2016). The levels of sphingolipid pathway metabolites, including ceramides and sphingosine-1-phosphate (S1P), are significantly deregulated in breast tumor tissue compared to normal tissue (Moro et al., 2018; Nagahashi et al., 2016), and elevated tumor ceramide levels have been shown to be associated with higher tumor grades (Moro et al., 2018; Schiffmann et al., 2009). The conversion of ceramides to sphingosine and subsequent phosphorylation generates S1P. The balance between the intracellular levels of pro-apoptotic ceramide and pro-survival S1P determines the fate of cancer cells (Spiegel and Milstien, 2002). By mediating apoptosis, growth arrest and senescence, ceramides function as tumor-suppressor lipids, whereas S1P is a key tumor-promoting lipid that enhances cell proliferation, migration and angiogenesis (Brocklyn, 2010; Cuvillier et al., 1996; Zhang et al., 1991).

In this study, we identify PAQR4 as a Golgi-localized ceramidase that is highly expressed in breast cancer tissues and whose expression is required for tumor growth. We find that enhanced PAQR4 expression endows cancer cells with a dual selective advantage through the combined effects of lowering cytotoxic ceramides and generating S1P.

## Results

### The Sphingolipid Metabolism-Related Gene *PAQR4* is Required for Breast Cancer Cellular Growth and Negatively Correlated with Breast Cancer Patient Survival

Gene set enrichment analysis (GSEA) of differentially expressed genes between 111 paired tumor and non-tumor tissue revealed an enrichment of genes in the ceramide signaling pathway (NES = 1.67, FDR = 0.075) and sphingolipid metabolism (NES = 1.62, FDR = 0.053) in tumor tissues (Figure 1A, S1A). To uncover key regulatory members of the sphingolipid pathway, we next performed a supervised differential expression analysis of sphingolipid-related genes (203 genes) between matched tumor and non-tumor tissue samples. We identified PAQR4 as the most upregulated gene transcript (FC∼5, adj. p-value < 0.01; Figure 1B). PAQR4 deregulation was breast cancer subtype-independent, as enhanced PAQR4 expression was detected in tumors irrespective of estrogen, progesterone receptor and HER2 receptor profile (Figure S1B). Consistent with the transcriptomic analysis, detection of PAQR4 by immunohistochemistry in breast cancer tissue sections showed enhanced staining intensity in the tumor compartment compared to the stromal/normal tissue areas (Figure 1C). PAQR4 expression was further negatively correlated to patient relapse-free survival (Log-Rank p-value = 0.0082) and disease-specific survival (Log-Rank p-value = 0.0301) in an in-house data set (Chrisanthar et al., 2011) (Figure 1D), as well as metastasis-free survival in an independent validation set (Figure S1C). These findings establish that PAQR4 expression is induced in tumor tissues and is negatively correlated to patient survival.

**Figure 1:**
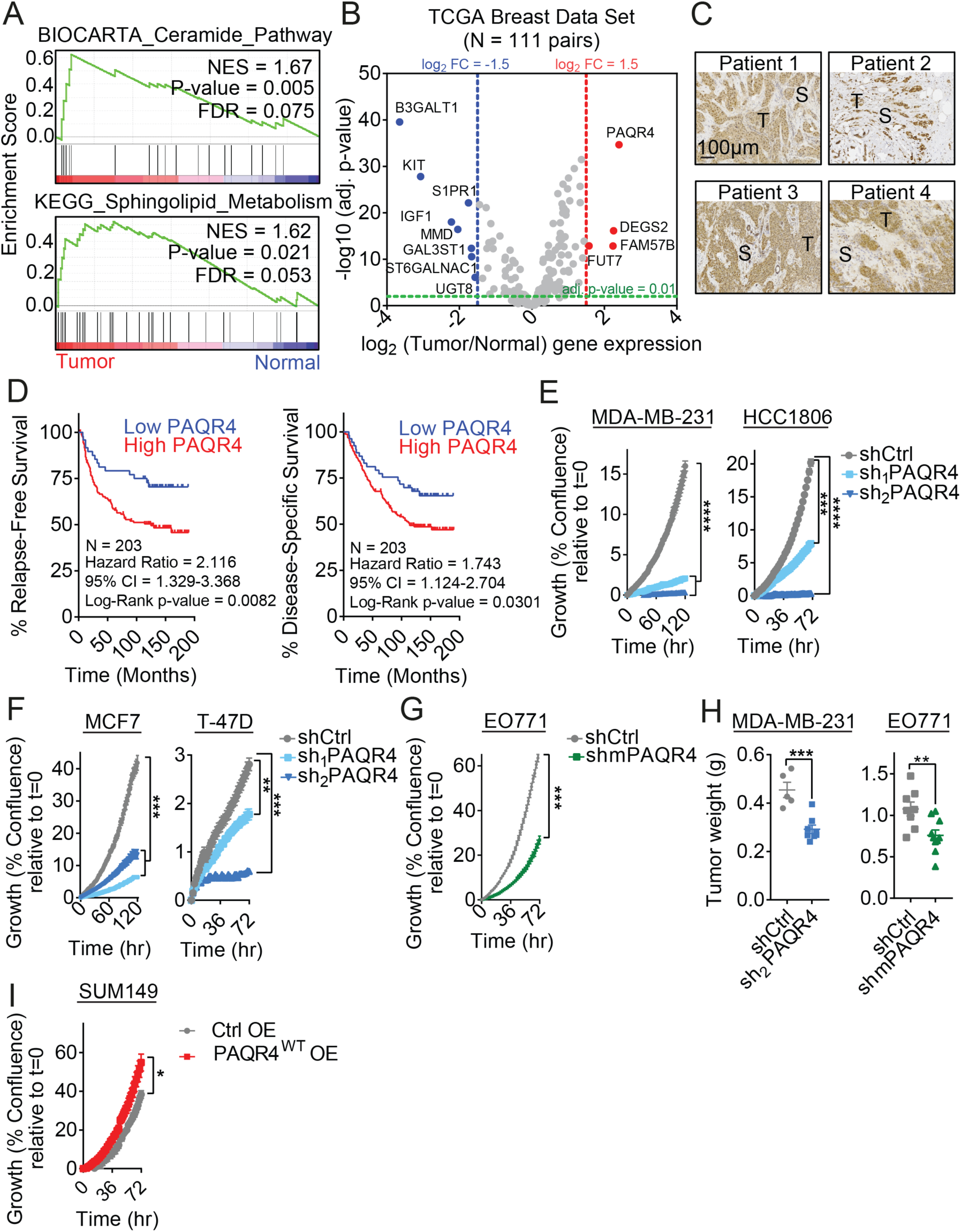
The Sphingolipid Metabolism-Related Gene *PAQR4* is Required for Breast Cancer Cellular Growth and Negatively Correlated with Breast Cancer Patient Survival. **A)** Functional analysis of differentially expressed genes comparing the transcriptomics of paired normal and tumor tissues from breast cancer patients (N=111) obtained from The Cancer Genome Atlas (TCGA) reveals an enrichment of genes in the ceramides signaling and sphingolipid pathway in tumor tissue. NES = Normalized Enrichment Score. FDR = False Discovery Rate. **B)** Volcano plot showing deregulated sphingolipid-related genes when comparing transcriptomic profiles of paired normal and tumor tissue derived from breast cancer patients (n=111). The significant (adjusted p-value < 0.01) over-expressed genes (red circles) are represented as log2 (tumor/normal) gene expression >1.5 and down-regulated (blue circles) as log2 (tumor/normal) gene expression < 1.5 fold changes. **C)** Representative PAQR4 staining in breast cancer patient tissue. S = stroma. T = tumor tissue. **D)** PAQR4 protein expression negatively correlates with patient survival (n=203) as determined by Kaplan-Meier curves of relapse-free and disease-specific survival based on high (equal or more than 25% quartile, red curve) or low (25% quartile, blue curve) PAQR4 expression using Log-rank test. **E, F, G)** Growth curve of triple negative (MDA-MB-231 and HCC1806), hormone receptor positive (MCF7 and T-47D) and murine EO771 cells transduced with lentiviral-delivered shRNAs targeting *PAQR4*. Cell growth was determined by high content imaging and represented as % confluence normalized to t=0, n≥6. Data are shown as mean±SEM. **H)** Tumor weight of orthotopically implanted MDA-MB-231 and EO771 cells with the indicated shRNAs targeting *PAQR4*. One representative of two experiments is shown as mean±SEM, n=4 mice (MDA-MB-231) or n=4-5 mice (EO771). **I)** Growth curve of SUM149 cells overexpressing PAQR4 in starved (1% FBS) medium. Cell growth was determined by high content imaging and represented as % confluence normalized to t=0, n=6. Data are shown as mean±SEM. \**P*<0.05, \*\**P*<0.01, \*\*\**P*<0.001

To further investigate the mechanism by which PAQR4 affects tumor cells, we next sought to determine the cellular phenotypes it governs. Depletion of endogenous PAQR4 in triple negative (MDA-MB-231 and HCC1806) and estrogen receptor positive (MCF7 and T47D) human breast cancer cell lines as well as the murine breast cancer cell line EO771, using lentiviral delivered short hairpin RNAs (shRNAs; Figure S1D) significantly reduced cellular growth as determined by high content imaging (Figure 1E, F and G). Such conserved effect across tumor cell genotypes is consistent with the observed PAQR4 induction between breast cancer subtypes. In contrast to the findings from Zhang, *et. al*. (Zhang et al., 2018), the observed reduction of cellular growth in PAQR4 depleted cells was independent of changes in cell cycle status (Figure S1E). Consistent with the diminished growth *in vitro*, PAQR4 depletion in human MDA-MB-231 and mouse EO771 triple negative breast cancer cells abrogated orthotopic tumor growth in immune-compromised NOD-SCID and immunocompetent C57BL/6J mice, respectively (Figure 1H). Finally, ectopic expression of PAQR4 in SUM149 cells increased the cellular growth (Fig. 1I, S1F). Collectively, these data establish that the sphingolipid metabolism-related gene PAQR4 is required for breast cancer cell growth.

### Homology Modelling Suggest that PAQR4 Functions as a Ceramidase

To investigate the mechanism by which PAQR4 regulates tumor growth, we next sought to determine the structural basis of its function. PAQR4 belongs to the PAQR protein family consisting of 11 members characterized by 7 transmembrane domains. Close family member PAQR2 (also known as AdipoR2) is known to possess ceramidase activity and has been co-crystalized with oleic acid - the byproduct of its enzymatic activity (Tanabe et al., 2015; Vasiliauskaite-Brooks et al., 2017). Within its catalytic site, a zinc ion, coordinated by three histidine residues (His202, His348 and His352), is essential for catalysis. Together with the zinc ion, an aspartic acid residue (Asp219) facilitates the cleavage of ceramide into sphingosine and a fatty acyl chain. Interestingly, these histidine residues and the aspartic acid are conserved in PAQR4 (His100, His240, His244 and Asp119; Figure S2A,B). We therefore constructed a homology model of PAQR4 based on the resolved crystal structure of AdipoR2. We then did a structural superimposition of the AdipoR2 structure to our PAQR4 model to position oleic acid inside the channel of PAQR4 and performed a 100 ns molecular dynamics simulation of the complex embedded in a lipid bilayer to relax the structure. This model revealed conservation of a 7-transmembrane structure that forms a barrel structure with an amphipathic pore (Figure 2A). The interactions of the fatty acid with PAQR4 were essentially the same as observed for AdipoR2. The carboxylic group of the fatty acyl locates at the proximity of the zinc coordination in PAQR4 (Figure 2A, B). We then repeated the simulation by docking a C18:1-ceramide molecule in the PAQR4 model using the oleic acid as an anchor. Independent simulations over 1.5 microseconds showed that PAQR4 - as was shown for PAQR2 (Vasiliauskaite-Brooks et al., 2017) - readily accommodated the fatty acyl part of the ceramide molecule inside the pore, with the cleavage site (amide bond) in close proximity to the active site zinc center (Figure 2C, Movie S2). Interestingly, in both structures, the sphingosine moiety of the ceramide molecule is predicted to localize outside of the pore. We further found that the homology model was remarkably stable over time (Figure 2D). To better understand the role of the zinc molecule we performed another simulation of C18:1-ceramide docked in PAQR4 wherein the three zinc-binding histidine residues were mutated to alanine – thereby excluding zinc from the structure. Interestingly, this simulation showed that both the fatty acid and the sphingosine arm of ceramide can be accommodated inside the amphipathic pore (Figure 2E). Although the overall structure of mutated PAQR4 were stable (Figure 2F), the more relaxed structure revealed that ceramide can interact in two different conformations. In sum, these modeling experiments suggest that PAQR4 possess ceramidase activity.

**Figure 2:**
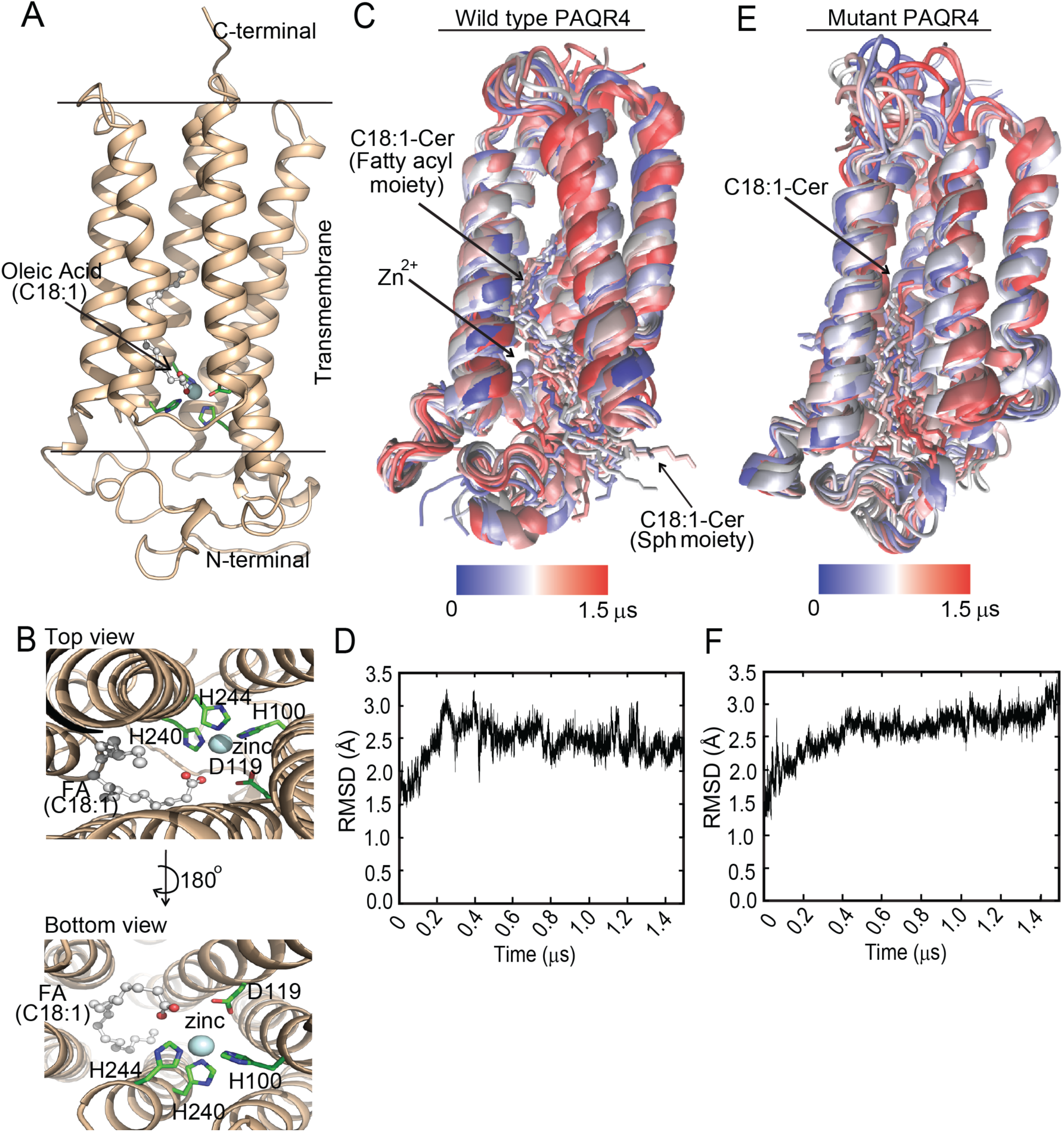
Homology Modelling Suggests that PAQR4 Function as a Ceramidase. **A, B)** Homology model of PAQR4 resolved within the membrane plane using PAQR2 as template structure. Top and bottom view of the hydrophobic binding pocket of PAQR4 is shown in B. Oleic acid (C18:1) in the amphipathic pore of PAQR4 is shown in ball and stick representation (with carbons in grey and oxygen in red). The three Zn-coordinated His together with Asp 219 are shown in stick representation. Zn is shown as a light blue sphere. **C)** Schematic molecular dynamic simulation of wild type PAQR4 in complex with C18:1-ceramide, embedded in a lipid bilayer. Frames of the receptor-ceramide complex, showing 1 frame every μs of the 1.5 μs simulation, color coded blue to red through the trajectory. Lipid molecules and water are omitted from the presentation for clarity. **D)** Root-mean-square deviation (RMSD) of the wild type PAQR4 protein atomic positions relative to first frame in the trajectory, as a function of simulation time. **E)** Schematic molecular dynamic simulation of mutant PAQR4 in complex with C18:1-ceramide, where the three Zn-coordinated histidines are mutated to alanine, embedded in a lipid bilayer. Frames of the mutated receptor-ceramide complex, showing 1 frame every μs of the 1.5 μs simulation, color coded blue to red through the trajectory. Lipid molecules and water are omitted from the presentation for clarity. **F)** Root-mean-square deviation (RMSD) of the mutant PAQR4 protein atomic positions relative to first frame in the trajectory, as a function of simulation time.

### PAQR4 is a Golgi-Associated Ceramidase

To experimentally test if PAQR4 harbors intrinsic ceramidase activity, we incubated deuterium-labeled ceramides with cellular lysates from control and PAQR4 knockdown cells. Following the incubation period, ceramidase activity was assessed by measuring the abundance of exogenous deuterated ceramide species by liquid chromatography-tandem mass spectrometry (LC-MS/MS). Consistently, we found that lysates from PAQR4 depleted cells displayed reduced ceramidase activity to that of the control cells (Figure 3A). We did not detect any PAQR4-related biases for specific ceramide fatty acyl chain length, as the levels of both ceramide species with long (C16:0 and C18:0) and very long fatty acyl chains (C24:1 and C24:0) were increased in knockdown lysates (Figure 3A). Given that we found PAQR4 to be induced in breast cancer, we wondered if this was paralleled by higher ceramidase activity in tumor compared to non-tumor tissue. To test this, we performed the ceramidase assay in matched normal and breast cancer tissue samples. A principle component analysis was able to differentiate tumor tissues from the corresponding normal tissues (Figure S3A). Interestingly, we found that tumor tissue lysates exhibited increased ceramidase activity for C16:0, C18:0 and C24:1 ceramides compared to their paired normal tissues lysates (Figure 3B). However, C24:0 ceramide abundance was unchanged. In further support of increased ceramidase activity in tumors, we found significant increase of labeled sphingosine – the downstream product of the ceramidase reaction – in the tumor compared to the matched control tissues (Figure 3C). Combined, these studies strongly suggest that PAQR4 possesses ceramidase activity. In addition to the aforementioned PAQR2, cancer cells contain several other ceramidases. A transcriptional analysis of these ceramidases revealed that PAQR4 is induced to a larger extent compared to other ceramidases (Figure S3B).

**Figure 3:**
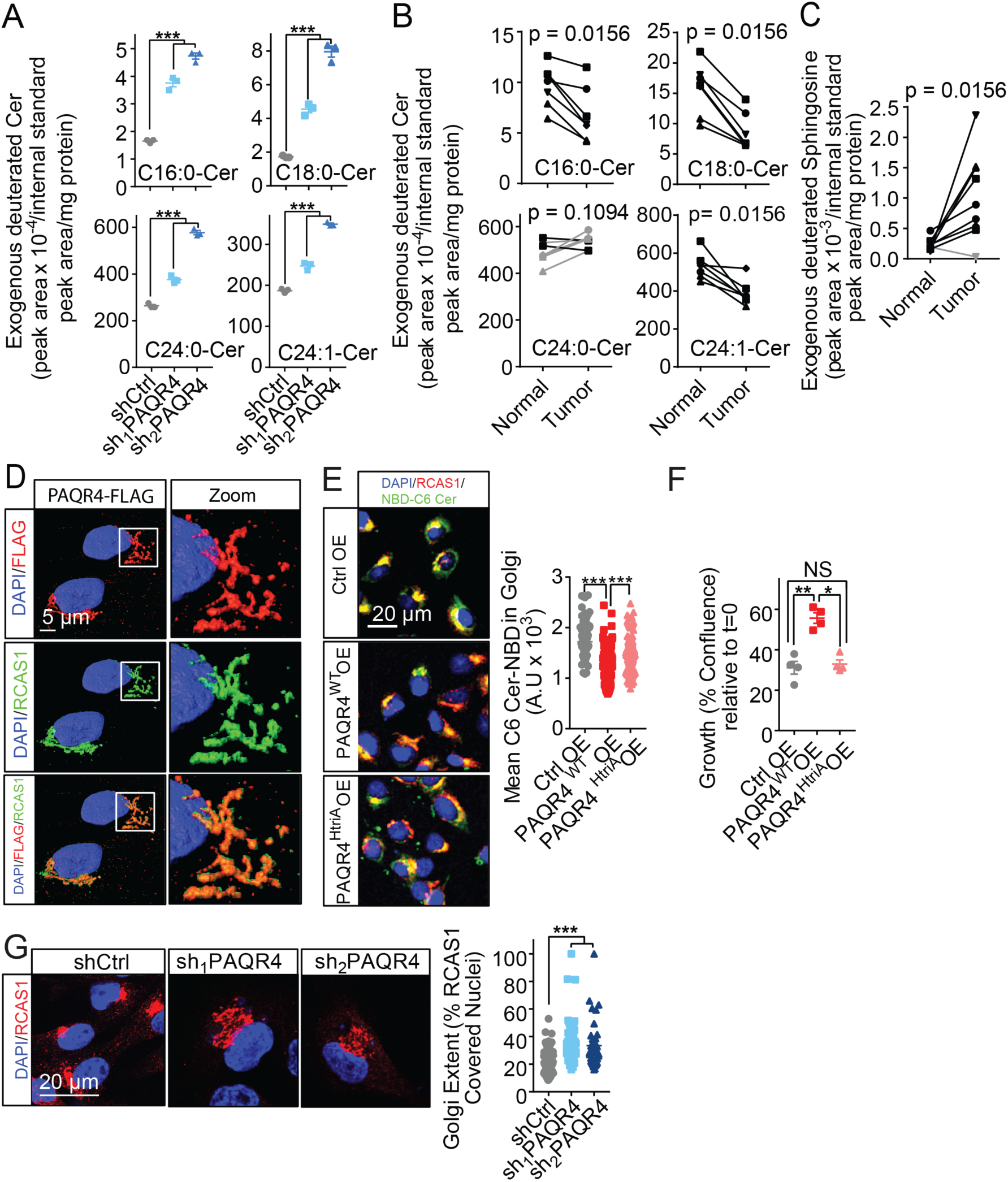
PAQR4 is a Golgi-associated ceramidase. **A)** PAQR4 ceramidase activity determined by incubating cell lysates from control and PAQR4 depleted cells with isotope (deuterium) labeled ceramides. Ceramide content was determined by LC-MS/MS and normalized to internal standard peak area and mg protein. One of two independent experiments is shown. Dot plot shows mean±SEM (n=3 per group). **B)** Ceramidase activity in paired breast cancer and normal tissue (n=7 patients) determined as described in A. Black lines represent patients with increased ceramidase activity in the tumor compared to normal tissue, grey lines represent patients with reduced activity in the tumor tissue. **C)** In addition to the levels of deuterium-labeled ceramides, the cleavage product, deuterium-labeled sphingosine, was determined by LC-MS/MS in the same lysates as in B. **D)** 3D confocal immunofluorescent images of MCF10A cells expressing Flag-tagged PAQR4. The tagged protein co-localizes with the Golgi marker RCAS1. Presented images were made by using the surface-rendering tool in the Imaris 9.1.2 Bitplane software. **E)** Ceramidase activity of PAQR4 in the Golgi determined by quantifying NBD-C6 ceramide fluorescence intensity in the Golgi (RCAS1-positive area) in SUM149 cells expressing WT or mutated PAQR4. The graph shows mean±SEM from one of three independent experiments (n≥30 cells/condition). **F)** Growth of SUM149 cells overexpressing WT or mutated PAQR4 incubated in starved (1% FBS) medium for 72 hrs. Cell growth was determined by high content imaging and represented as % confluence normalized to t=0, n=4. Data are shown as mean±SEM. **G)** Confocal immunofluorescent images of Golgi structure in MDA-MB-231 cells depleted for PAQR4. Golgi extent was measured as % RCAS1-covered peri-nuclei. Data are presented as mean ±SEM from one of three independent experiments (n≥30 cells/condition). * *P*<0.05, \*\**P*<0.01, \*\*\**P*<0.001.

Sphingolipid metabolism is highly compartmentalized within the cell (Hannun and Obeid, 2008); thus, we next used 3D confocal imaging to determine the subcellular localization of PAQR4. Detection of expressed flag-tagged PAQR4 revealed a peri-nuclear staining that co-localized with the Golgi marker RCAS1 both in MCF10A and breast cancer cell line SUM149 (Figure 3D, S3C, S3D). To further determine if PAQR4 possesses ceramidase activity in the Golgi compartment, we took advantage of the fact that NBD fluorescent-labeled C6 ceramide readily accumulates in the Golgi when added exogenously to cells (Lipsky and Pagano, 1985; Pagano et al., 1989). We therefore added NBD-labeled ceramides to control and PAQR4 overexpressing cells and measured Golgi-localized fluorescence intensities by confocal imaging. Interestingly, we found that overexpression of PAQR4 significantly reduced the NBD-labeled ceramide in the Golgi as determined by co-signal between NBD and RCAS1 (Figure 3E). Further, this effect was significantly reduced when the three histidine residues coordinated with the zinc ion in the catalytic domain were mutated to alanine (PAQR4^HtriA^OE; Figure S3E). At the cellular level, mutation of the three histidine residues, ablated the ability of PAQR4 to promote cancer growth (Figure 3F).

Ceramides accumulation in the Golgi has been shown to induce local Golgi fragmentation (Sakamoto et al., 2018). Consistent with PAQR4 having ceramidase activity, we observed a significant Golgi fragmentation in the PAQR4 depleted cells compared to the control cells by confocal immunofluorescence (Figure 3G). Collectively, this suggests PAQR4 acts as a ceramidase in the Golgi compartment.

### PAQR4 Depletion Causes Accumulation of *de novo* Sphingolipid Intermediates and Ceramide-induced Apoptosis

Having identified PAQR4 as a ceramidase in the Golgi, we next sought to determine how PAQR4 depletion alters cellular sphingolipid homeostasis. To this end, we performed an unbiased sphingolipidomics analysis of both MDA-MB-231 and MCF7 cells with depleted levels of PAQR4. Consistent with a ceramidase activity, we found overall elevated levels of ceramides in the PAQR4 knockdown cells in both cell lines (Figure 4A). Similar to what we observed for ceramidase activity (Figure 3A and 3E), we did not detect any PAQR4-related biases for specific ceramide fatty acyl chain length. Interestingly, we also detected an accumulation of upstream intermediates (i.e. dihydroceramides and dihydrosphingosine) in the *de novo* sphingolipid synthesis pathway (Figure S4A, 4B and 4C), but not the downstream metabolites (i.e. glycosylceramides and sphingomyelins) (Figure S4B and S4C). This suggests that the *de novo* synthesis pathway is highly active in breast cancer cells. Accumulation of cellular ceramides is tightly linked to apoptosis (Jarvis et al., 1996; Obeid et al., 1993). We therefore asked if the reduced growth rates observed in the PAQR4 depleted cells is caused by increased apoptosis by assessing externalized phosphatidylserine in MDA-MB-231, HCC1806, MCF7 and T-47D PAQR4 knockdown cells by flow cytometry. Accordingly, we found increased rates of apoptosis across all cell lines (Figure 4D). Further, immuno-staining of cleaved caspase-3 in tumor sections from control and PAQR4 knockdown tumors grown *in vivo*, confirmed activation of the apoptotic pathway in PAQR4 depleted cells (Figure 4E). Ceramides are linked to apoptosis by forming pores in the outer mitochondrial membrane to facilitate the release of cytochrome C from mitochondria into the cytoplasm (Siskind et al., 2006; Tomassini and Testi, 2002). Consistent with our hypothesis and cellular buildup of ceramides, we found increased levels of cytoplasmic cytochrome C in PAQR4 depleted cells (Figure 4F). Considering the selective advantage of PAQR4 induction in tumors, we postulated that in addition to reducing cytotoxic ceramides, induction of PAQR4 could provide intermediates for S1P production. In support of this, we found that PAQR4 co-localizes with the downstream kinase, sphingosine kinase-1 (SPHK1), in the Golgi compartment (Figure 4G). Further, overexpression of wild type PAQR4 increased cellular S1P levels, while overexpression of the ceramidase-dead HtriA mutation did not (Figure 4H). In sum, we find that PAQR4 depletion results in ceramide-induced apoptosis and an accumulation of the *de novo* sphingolipid synthesis intermediates.

**Figure 4:**
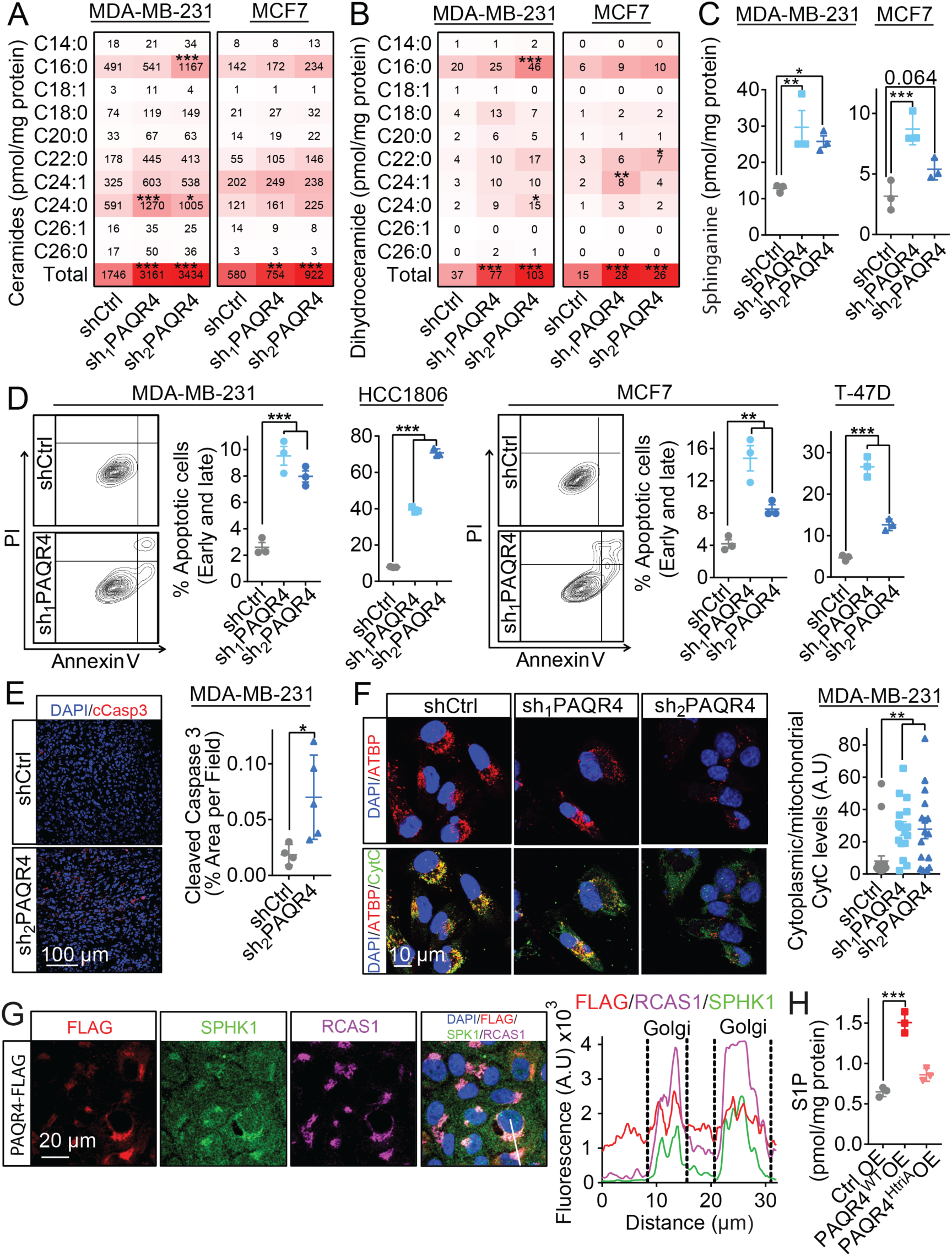
PAQR4 Depletion Causes Accumulation of *de novo* Sphingolipid Intermediates and Ceramide-induced Apoptosis. **A, B)** Ceramides and dihydroceramides levels in lysates from MDA-MB-231 and MCF7 cells depleted for PAQR4 determined by liquid chromatography-tandem mass spectrometry (LC MS/MS). Sphingolipid levels were normalized to the protein concentration and are presented as mean in the heatmap, n=3 per group from one of two independent experiments. **C)** Sphinganine levels in lysates from MDA-MB-231 and MCF7 cells depleted for PAQR4 determined as in A and B. The sphingolipid levels were normalized to the protein concentration and are presented as dot plots in mean±SEM, n= 3 from one of two independent experiments. **D)** Percentage of apoptotic cells (early and late) upon loss of PAQR4 in MDA-MB-231, HCC1806, MCF7 and T-47D cells determined by AnnexinV and PI staining measured by flow cytometry. Dot plots show mean±SEM, n= 3 from one of more than three individual experiments. **E)** Confocal immunofluorescent staining of cleaved caspase 3 (cCasp3) as an apoptosis marker in tumor sections of MDA-MB-231 tumors lacking PAQR4 expression. cCasp3 staining was quantified as %cCasp3 positive area per field and data are shown in dot plot as mean±SEM, n= 5/group. **F)** Immunofluorescent staining of cytochrome C as an indicator of ceramide-induced apoptosis in PAQR4 depleted MDA-MB-231 cells. Cytochrome C levels were quantified in the cytoplasm and in the mitochondria (ATPB positive area). Ratios are presented in the dot plot, which shows mean±SEM. (n= 30 cells) from one of three individual experiments. **G)** Immunofluorescent images of sphingosine kinase 1 (SPHK1) localization in MCF10A cells expressing Flag-tagged PAQR4. On the right side, co-localization of PAQR4 and SPHK1 in the Golgi (RCAS1) is illustrated with a line plot of fluorescence intensity, based on the white line shown in the rightmost merged image. **H)** S1P levels in lysates from MDA-MB-231 cells with PAQR4 overexpression was measured by LC MS/MS. S1P levels were normalized to protein concentration and are presented as mean, n=3 per group. One of two independent experiments is shown. \**P*< 0.05, \*\**P*<0.01, \*\*\**P*<0.001.

## Discussion

Beyond their structural role in the membrane homeostasis, sphingolipids control critical signaling transduction pathways within the cancer cells to drive growth, proliferation, migration and invasion. Altered levels of ceramide species and other metabolites in the sphingolipid metabolism network of the cancer cells is related to the deregulation of the expression of the genes encoding the enzymes within this network (Moro et al., 2018). Here, we identify and molecularly define the role of the ceramidase PAQR4 in breast tumors. We find that PAQR4 is highly expressed in the tumor tissue of the breast cancer patients compared their corresponding normal tissue and its expression negatively correlates to patient survival. Based on our findings, we propose that PAQR4 acts as a Golgi-localized ceramidase that provides a selective advantage to cancer cells through reducing cytotoxic ceramides as well as providing the building blocks for S1P production. Consistent with the work by Zhang *et. al.* (Wang et al., 2017; Wu and Liu, 2019; Zhang et al., 2018), we find that PAQR4 reduces cancer cell growth. Their studies suggest that PAQR4 controls cellular growth through stabilization of cyclin-dependent kinase 4 (CDK4) and hence cell cycle state. We were not able to experimentally observe cell cycle arrests in PAQR4 depleted cells. Instead, through a combination of structural homology modeling, enzymatic and lipidomics approaches, we propose that PAQR4 functions as a ceramidase. As a consequence of reduced ceramidase activity, PAQR4 knockdown cells accumulate ceramides – a lipid shown to regulate CDK4 levels (Pastukhov et al., 2014). The *de novo* synthesis of ceramides is initiated in the endoplasmic reticulum by condensation of serine and fatty acids by the serine palmitoyltransferase complex (Braun and Snell, 1967). Ceramides are then converted enzymatically into different classes of sphingolipids in the ER, cis- and medial-Golgi network that subsequently are integrated in organelle membranes (Ogretmen, 2018). In this study, we show that PAQR4 is localized to the Golgi apparatus where it degrades ceramides into sphingosine, which could be further phosphorylated into S1P by the co-localized SPHK1. Although PAQR4 is homologous with other PAQR family members – in particular PAQR1-3, PAQR4 appear to be selectively induced in breast cancer tissue compared to patient-matched non-tumor tissue. We speculate that the selective advantage of PAQR4 could be related to its placement in the Golgi. Other members, PAQR1 and 2 (commonly known as AdipoR1 and 2), are placed in the plasma membrane, where they serve as receptors for the anti-diabetic hormone adiponectin (Yamauchi et al., 2003; Yamauchi et al., 2007). By intervening early in the sphingolipid pathway, we suggest that the tumor cells gain a selective advantage by being able to reduce apoptotic ceramide species while at the same time create the intermediates required for S1P synthesis. We did not detect PAQR4 in the plasma membrane, but it is tempting to speculate if PAQR4 activity could be regulated by an intracellular ligand.

Finally, despite the local placement in the Golgi apparatus, we find evidence of global cellular accumulation of ceramides. In particular, the release of mitochondrial cytochrome C to the cytoplasm strongly suggests a mitochondrial buildup of ceramides. Future work will determine if this is a result of direct Golgi-mitochondrial interaction sites (Dolman et al., 2005) or vesicular lipid transfer (Voelker, 1990).

Here we describe how PAQR4 provides cancer cells with an adaptive advantage for growth through modulation of the balance between ceramides and S1P. This suggests that development of small molecule inhibitors of PAQR4 could provide a feasible path for a novel class of targeted therapies for breast cancer either as a stand-alone treatment or in combination with chemotherapies known to alter ceramide levels.

## ACKNOWLEDGMENTS

We thank James Lorens and Claudio Alacórn for comments on previous versions of the manuscript. We thank the VCU Lipidomics/Metabolomics Core, the NIH-NCI Cancer Center Support Grant P30 CA016059 to the VCU Massey Cancer Center, as well as a shared resource grant (S10RR031535) from the National Institutes of Health for assistance with lipidomics analysis. We also acknowledge the Flow Cytometry Core Facility, Department of Clinical Science, and the Molecular Imaging center, Department of Biomedicine, University of Bergen. N.H. is supported by the Trond Mohn Foundation Starting Grant.

## AUTHOR CONTRIBUTIONS

N.H conceived the project and supervised all the research. L.P, P.P and N.H wrote the manuscript. L.P, P.P and M.I.S designed, performed and analyzed the experiments. R.G and P.E.S developed and performed the ceramidase assays. S.K, P.E.L and A.M collected and provided patient samples. K.T. performed structural homology analysis and molecular dynamics simulations.

## STAR methods

### Lead Contact and Materials Availability

Further information and requests for resources and reagents should be directed to and will be fulfilled by the Lead Contact, Nils Halberg. This study did not generate new unique reagents.

### Experimental Model and Subject Details

#### Cells

Cells were maintained using standard tissue culture procedures and grown at 37°C with 5% CO_2_ and atmospheric oxygen. MDA-MB-231 (human, female), EO771 (mouse, female) and HEK293T (human, fetus) cells were cultured in Dulbecco’s modified Eagles medium (DMEM, Sigma, D5671) supplemented with 10% fetal bovine serum (FBS, Sigma, F7524) and 2mM L-glutamine (Sigma, G7513). HCC1806 (human, female) cells were cultured in RPMI-1640 medium (Sigma, R8758) supplemented with 10% FBS. MCF7 (human, female) cells were cultured in Minimum Essential medium (MEM, Sigma, M4655) supplemented with 10% FBS and 1% MEM Non-essential amino acids solution (NEAA, Sigma, M7145). T-47D cells were cultured in RPMI-1640 medium supplemented with 10% FBS and 10 μg/ml human insulin solution (Sigma, I9278). MCF10A (human, female) were cultured in DMEM/F12 supplemented with 5% horse serum (Sigma, H1270), 20 ng/ml human epidermal growth factor (hEGF, Sigma, E9644), 0.5 mg/ml hydrocortisone (Sigma, H6909), 100 ng/ml cholera toxin (Sigma, C8052), and 10μg/ml human insulin. SUM149 cells were cultured in DMEM/F12 medium supplemented with 5% FBS, 1 mg/ml hydrocortisone and 5μg/ml human insulin. All media contained 100 U/ml penicillin and 100ug/ml streptomycin. Ishikawa cells (human, female) were cultured in MEM supplemented with 5% FBS and 1% NEAA.

For cell line authentication, MDA231, MCF7 and MCF10A cells were harvested for genomic DNA extraction using Genomic DNA isolation kit (Norgen Biotek, 24700). Isolated genomic DNA was analyzed by Eurofins Genomics laboratory and the cell lines authenticated based on genetic fingerprinting and short tandem repeat (STR) profiling.

#### Patient Tumor Specimens

Female breast cancer patient specimens used for ceramidase assay (Figure 3B,C), were obtained at the Department of Surgery (Haukeland University Hospital, Bergen, Norway). Tumor tissue and matching benign tissue from a non-tumor bearing quadrant of the breast was obtained from mastectomy specimens collected as described in a previously (Lonning et al., 2009). Breast cancer tissue sections used in Figure 1C were from anonymized patients included in the study by Chrisanthar *et. al.* (Chrisanthar et al., 2011).

#### Animal Models

Animal experiments were approved by the Norwegian Animal Research Authority and conducted according to the European Convention for the Protection of Vertebrates Used for Scientific Purposes, Norway. The Animal Care and Use Programs at University of Bergen are accredited by AAALAC international. The laboratory animal facility at University of Bergen was used for the housing and care of all mice.

Female NOD-SCID gamma mice were purchased from Jackson laboratories and maintained under defined flora conditions in individually ventilated (HEPA-filtered air) sterile micro-isolator cages (Tecniplast, Buguggiate, Italy). Bedding and cages were autoclaved and changed twice per month. C57BL/6J mice were obtained from Jackson Laboratories and bred on site. Mice were kept in IVC-II cages (Sealsafe® IVC Blue Line 1284L, Tecniplast, Buguggiate, Italy). For both strains, 5-6 mice were housed together and maintained under standard housing conditions at 21°C ± 0.5°C, 55% ± 5% humidity, and 12h artificial light-dark cycle (150 lux). Mice were provided with standard rodent chow (Special Diet Services RM1, 801151, Scanbur BK, Oslo Norway) and water *ab libitium*.

## METHOD DETAILS

### Syngeneic and Xenograft Model

For *in vivo* studies of PAQR4 depletion, cells were orthotopically implanted in the 4^th^ inguinal mammary fat pad of 8 (C57BL/6J) or 6 (NOD-SCID) weeks old female mice. Mice were anesthetized by isoflurane (2 mg/kg) and a small incision was made to visualize the mammary gland. 5×10^4^ viable EO771 or 5×10^5^ viable MDA-MB-231 cells in Phosphate Buffered Saline (PBS) were mixed 1:1 by volume with matrigel (Corning, 356231) and injected in a total volume of 50 μl. Mice were injected in both the left and right mammary fat pad. The incision was closed with wound clips (EZ Clips^TM^, 59027). At endpoint the mice were humanely euthanized and tumors harvested for weight measurements and histology.

### RNAseq Analysis Quantification

Total Transcriptomics profiles of paired normal and tumor tissues of 111 breast invasive carcinoma patients were obtained from The Cancer Genome Atlas (v1.5.2 TCGA). To identify the differentially expressed genes (DEGs), a supervised paired sample t-test were implemented using limma Bioconductor package in R (Ritchie et al., 2015). To investigate the DEGs involved in sphingolipid metabolism, we compiled a gene set containing 203 genes involved in the sphingolipid and ceramide biosynthesis, catabolism, and signaling pathways based on the Molecular Signatures Database (v6.1 MsigDB) (Subramanian et al., 2005). Adiponectin Receptor 1 and 2 (AdipoR1 and AdipoR2) are known as ceramidases. All 11 genes in the progestin and adipoQ receptor (PAQR) family were also included in the gene set. Adjusted p-value 0.01 and log2 tumor/normal fold change expression ±1.5 were used as cut-offs to select differentially expressed genes.

### Gene Set Enrichment Analysis (GSEA)

Functional Analysis of the DEGs comparing the transcriptomics of the paired normal and tumor tissues from TCGA was performed using GSEA software (Subramanian et al., 2005). The significant enriched gene sets were selected as FDR < 0.25.

### Survival Analysis

Survival analysis was performed on a quartile normalized cDNA microarray data of breast patient’s tumor samples based on Illumina ® HumanHT-12v4 Expression platform. The samples were treatment naïve and collected in a clinical trial assessing mechanisms of resistance to neoadjuvant chemotherapy locally advanced breast cancer (Chrisanthar et al., 2011). Disease-Specific (DSS) and Relapse-Free Survival (RFS) were calculated with GraphPad Prism software using Log-rank test. Patients with 25% quartile of PAQR4 expression were considered as low PAQR4 group and patients with equal and more than 25% quartile were classified as patients with highly expressed PAQR4 tumors. In addition, Gene expression-based Outcome for Breast cancer Online (GOBO) (v.1.0.3) (Ringner et al., 2011) were used to graph the distant metastasis-free survival based on PAQR4 expression in 1176 available breast tumor.

### Structural Prediction and Molecular Dynamic Simulation

PAQR2 and PAQR4 amino acid sequence alignment were performed using Clustal omega. Swiss-Model (Waterhouse et al., 2018) was used to prepare a homology-model of PAQR4, based on the structure of AdipoR2 (PDB ID: 5LXA). The PAQR4 model was embedded in a lipid bilayer using the Charmm membrane builder (Wu et al., 2014) and subjected to molecular dynamics simulations with the Amber molecular modelling package as described below.

Simulations of PAQR4 in complex with ceramide were based on docking of ceramide to the homology model of PAQR4. The PAQR4-ceramide complex was embedded in an equilibrated bilayer of POPC (Palmitoyl-oleyl-sn-glycero-3-phosphocholine). The system was solvated in a water box encompassing the protein with extensions in the XY-dimensions identical to the lipid bilayer. Starting coordinates for simulations of PAQR4 with the three zinc-coordinating histidine residues mutated to alanine were prepared by removing zinc and modifying histidine to alanine in the coordinate file. Simulations were performed using the GPU-accelerated PMEMD module (Salomon-Ferrer et al., 2013) implemented in the AMBER18 molecular dynamics software package (D.A. Case, 2016), applying the ff14SB (Maier et al., 2015) and lipid14 force fields (Dickson et al., 2014). Periodic boundary conditions with particle mesh Ewald summation (Darden et al., 1993) of electrostatic interactions were applied and van der Waals interactions were truncated with a 10Å-cutoff. SHAKE (Elber et al., 2011) was used to constrain bonds involving hydrogen atoms. After 10,000 steps of minimization, the systems were subjected to 5 ps of gradual constant volume heating (NVT) from 0 to 100 K followed by 100 ps gradual constant pressure heating (NPT) applying anisotropic Berendsen coupling (Berendsen et al., 1984) with reference pressure set to 1 bar. Temperature was regulated with the Langewin thermostat (Loncharich et al., 1992) gradually over 100ps from 100 K to 303 K. Weak restraints maintained by a force constant of 10 kcal mol^-1^ Å^-2^ were applied to the protein and lipids throughout both heating steps. The systems were finally simulated at constant pressure conditions (NPT) without restraints for 1500 ns at 303K and the resulting coordinates were saved every 100 ps for analysis.

### Cloning of PAQR4

Full-length PAQR4 (Isoform 1) was PCR amplified from cDNA of Ishikawa cells and cloned into the retroviral pBabe-puro vector (Addgene #1764) by conventional restriction enzyme-based cloning. Sanger DNA sequencing analysis confirmed successful cloning.

### Mutagenesis

Site directed mutagenesis was performed on PAQR4 constructs by using the PCR-based QuickChange Lightning Multi Site-Directed Mutagenesis Kit (Agilent Technologies, 210515). Primers were designed by the use of Aligent’s primer design tool and mutagenesis was performed according to manufactures instructions. Primers are listed in the key resource table. Sanger sequencing analysis was performed to confirm mutated PAQR4 products.

### Knockdown and Overexpression Cell Lines

Short hairpin RNA (shRNA) constructs for PAQR4 and scramble (shCtrl) were purchased from Sigma (key resource table).

For generation of virus, HEK293T cells were plated in 10 cm plates to reach a confluency of 70% on the following day. For lentiviral production of shRNAs targeting PAQR4, 12 μg plasmid was co-transfected with 6 μg packing plasmid and 12 μg VSVG envelope plasmid. pBabe-puro vector containing PAQR4 was co-transfected into HEK293T cells with the retroviral envelope VSVG (12 μg) and packaging Gag/Pol (6 μg) plasmids to produce retroviral particles. Transfections were performed in antibiotic-free media using 60 μl Lipofetamine2000 according to manufactures protocol. After 16 hours, media was replaced with fresh media containing penicillin and streptomycin. 48 hours after transfection, virus-containing supernatant was harvested and spun at 290 g before being filtered through a 0.45-μm filter. Next, virus was used to infect sub-confluent cultures in the presence of polybrene (10 μg/ml) overnight. 48 hours after infection, antibiotic selection was performed with the following concentrations of puromycin; 4 μg/ml for MCF10A, 2 μg/ml for MDA-MB-231, T47-D and EO771 and 1.33 μg/ml for MCF7 and HCC1806. Uninfected control cells were kept alongside to determine the endpoint of selection (2-3 days). After selection, cells were allowed to recover for at least 48h before tested for overexpression or knockdown of PAQR4. Knockdown cells were never passaged for more than 3 passages after infection.

### RT-qPCR

The efficiency of PAQR4 knockdown and overexpression was confirmed by RT-qPCR. Total RNA extraction was performed using total RNA purification kit (Norgen Biotek, 37500) according to the manufactureŕs protocol. cDNA synthesis was performed using Superscript III reverse transcriptase (Invitrogen, 18080051) with 1 μg RNA as template. Quantitative RT-PCR was performed using SYBR Green (Roche, 4707516001) on the LC480 instrument (Roche) according to standard procedures. PCR reactions were performed in quadruplicates and the relative amount of cDNA was calculated by the ΔΔCT method using human Hypoxanthine-guanine phosphoribosyltransferase (h*HPRT*) RNA expression as a control for human cell lines and mouse actin (m*Actin*) for the mouse cell line. Primers can be found in the resource table.

### Proliferation Assay

For cell proliferation assays 1.5 × 10^3^ MDA-MB-231, 1.5 × 10^3^ HCC1806, 1×10^3^ MCF7, 2.0 × 10^3^ T-47D and 5×10^2^ EO771 cells were seeded in 100 μl total volume per well in 96 well plates. Plates were centrifuged at 290g for 5 min, and cells were allowed to adhere overnight. The next day, 100 μl additional media were added before growth was followed using the IncuCyte Zoom (Essen Bioscience). For cell proliferation assays in SUM149, 1.1 × 10^3^ SUM149 cells were seeded out in 100 μl total volume per well in 96 well plates and the next day, the medium was substituted with 200 μl media containing 1% FBS. In all experiments cells were seeded in at least triplicates, and four fields were imaged per well under 10x magnification every 2 h for 3-5 days. The IncuCyte Zoom (v2018A) software was used to calculate confluency values. All proliferation assays were performed in triplicate.

### Tissue Immunohistochemistry and Immunofluorescence

For tissue immunohistochemistry and immunofluorescene, 5 µm tissue sections were deparaffinized in xylene (twice 10 min each), and rehydrated sequentially in ethanol (twice 10 min each in 100%, 10 min in 95%, 10 min in 75%) and twice, 5 min each, in distilled H_2_O. For immunohistochemistry of endogenous PAQR4 staining, antigen retrieval was performed by boiling the deparaffinized tissue slides for 20 min at 85-90°C in Tris-based antigen unmasking solution, pH 9.0 (Vector Laboratories, H-3301) and cooled down in 0.05% Tween20/PBS (washing buffer). Next, sections were blocked for 20 min in 1.5% goat serum provided by Vectastain ABC horseradish Peroxidase (HRP) kit (Vector Laboratories, PK-4001), and incubated overnight at 4°C with PAQR4 primary antibody diluted in 1% Bovine Serum Albumin (BSA) in PBS. Specimens were then washed three times for 10 min each in the washing buffer and incubated with BLOXALL endogenous peroxidase and alkaline phosphatase blocking solution (Vector Laboratories, SP-6000) for 10 min at room temperature (RT) and rinsed three times for 10 min each in washing buffer. Next, slides were incubated 30 min at RT with 0.5% biotinylated anti-rabbit secondary antibody from Vectastain ABC HRP kit. After rinsing three times for 10 min for each in the washing buffer, slides were treated 30 min at RT with Vectastain ABC reagent. The HRP signal was then blotted by incubating the sections with HRP substrate (Vector Laboratories, SK-4105) for 1 min at RT. Counterstaining was performed by immersing the slides for 30 sec in haematoxylin, followed by 10 min rinsing in running tap water. The sections were dehydrated sequentially in ethanol (3 min in 70%, twice 3 min each in 96%, and 2 min in 100%), and twice 2 min each in xylene, before mounted with Eukitt mounting media (Sigma, 03989). The images were acquired using Hamamatsu slide scanner and Aperio ImageScope software.

For immunofluorescence of Cleaved Caspase 3, mouse tumor tissue was fixed in 4% paraformaldehyde (PFA) overnight and embedded in paraffin for sectioning. Sections were cut at 5 μm. After deparaffinization and rehydration as described above, antigen unmasking was performed in sodium citrate buffer, pH 6.0 (Vector Laboratories, H-3300) at 95-98°C for 20 min. After cooling down sections at RT for 30 min, sections were washed twice in washing buffer and blocked in 4% normal goat serum in PBS with 1% BSA. Next, sections were stained with Cleaved Caspase 3 antibody, followed by incubation with secondary antibody (Alexa Fluor 555, Invitrogen). Finally, sections were counterstained with 1 μg/ml DAPI, mounted using Prolong Diamond antifade reagent (Invitrogen, P36970) and imaged using the Leica SP5 confocal microscope. Cleaved caspase 3 staining was quantified as stained area per field (∼0.15 mm^2^) in 5 different fields per tumor. n=5 per group.

### Cell Immunofluorescence and Confocal Microscopy

Cultured cells were seeded on round glass slides in 6-well dishes. Two days later, cells were washed once in PBS, fixed in 4% PFA for 10 min, washed three times in PBS and permeabilized with PBS + 0.1% Triton-X for 5 min. After three washes in PBS, non-specific binding was blocked by incubation in 5% goat and/or donkey serum for 30 min at RT. Cells were then stained with primary antibodies for 1 hour at RT, washed three times with PBS, and incubated with secondary antibodies for 1 hour at RT. Nuclei were stained with 1 μg/ml DAPI (Sigma, D9542) in PBS for 5 min followed by three washes in PBS. Stained cells were mounted on glass using Prolong Diamond antifade reagent and left to dry at RT overnight in the dark. The type, source and dilution of antibodies are described in the key resource table. Immunostained samples were imaged using Leica SP5 confocal microscope. Images for PAQR4 localization were acquired using Leica SP8 confocal microscope. Image analysis was performed using ImageJ Fiji. The surface-rendering tool in the Imaris 9.1.2 Bitplane software was used to generate Figure 3D.

### Flow Cytometry

#### Apoptosis

2×10^5^ MDA-MB-231, HCC1806 and MCF7 and 3 × 10^5^ T47-D cells were seeded out in triplicates in 6-well plates. To collect all live and apoptotic cells, both media and cells were harvested 3 days after plating. Cell suspensions were centrifuged at 805 g for 5 min and washed once with ice-cold PBS. Next, cells were resuspended in annexin binding buffer (10 mM HEPES, 140 mM NaCl, and 2.5 mM CaCl_2_, pH 7.4) and stained with Annexin V (ThermoFisher Scientific, A13201) and propidium iodide (PI, abcam, Ab139418) for 15 min at RT in dark according to the manufacturer’s instructions. Flow cytometry was performed using the Accuri C6 (BD Biosciences) and data were analyzed with FlowJo software (Tree Star, Inc).

#### Cell Cycle

For cell cycle assays, cells were plated at a density of 2×10^5^ cells in triplicates in 6-well plates, and harvested 3 days later. Cell suspensions were washed with PBS once and fixed with ice-cold 70% ethanol for 30 minutes on ice. Fixed cells were washed with PBS twice before adding PI/RNase Staining Buffer (BD BioSciences, 550825) according to the manufacturer’s instructions and analyzed by flow cytometry. Flow cytometry was performed using the Accuri C6 (BD Biosciences) and data were analyzed with FlowJo software (Tree Star, Inc).

#### Cellular Sphingolipidomics

MDA-MB-231 (2.5×10^5^) and MCF7 (2.5×10^5^) cells were seeded out in triplicates in 6 well plates and allowed to adhere overnight. The following day the media was renewed with fresh culture media. For cells subjected to S1P measurements the media was replaced with DMEM phenol free media (Sigma, D1145) containing 0.02% FBS. 2 days after plating, cells were washed once in PBS and harvested by scraping. Cell suspensions were spun at 300g for 5 min at 4°C, and pellets were snap-frozen. Sphingolipids were quantified by LC-MS/MS as described (Shaner et al., 2009) by the VCU Lipidomics/Metabolimics core. The measured sphingolipid levels were normalized to protein concentration as determined by Bicinchoninic Acid (BCA) assay using Pierce BCA Protein Assay Kit (Thermo Fischer Scientific, 23225).

### Ceramidase Activity Assay and Sphingolipid measurement

Tumor biopsies and frozen cell pellets were homogenized in 500 µl cold Dubelco’s PBS solution containing Calcium and Magnesium and EDTA-free protease inhibitor cocktail (Roche) with a mechanical tissue homogenizer (tissue biopsies) and tissue dismembrator probe (cell pellets, Fisher Scientific Model FB50, 8 short 1 seconds pulses at 15% of amplitude). The aqueous homogenate was transferred to a borosilicate glass tube. The original microcentrifuge tube was rinsed with additional 500 µl of cold Dubelco’s PBS solution and the aqueous solutions were combined. After incubation of the samples on ice, 10 µl of an ethanolic mixture of deuterated Ceramides-d7 was added to each sample (Ceramide d18:1-d7/16:0 (Cayman Chemicals, 22787, 4.9 µM), Ceramide d18:1-d7/18:0 (Cayman Chemicals, 22788, 4.6 µM), Ceramide d18:1-d7/24:1 (Avanti Polar Lipids, 10 µM), Ceramide d18:1-d7/24:0 (Avanti Polar Lipids, 860679, 10 µM). The homogenates were incubated at 37 °C with continuous shaking for three hours and the reaction was quenched by adding 2 ml of organic extraction solvent (Isopropanol/Ethyl Acetate 1:2, v:v). Immediately afterwards, 20 µl of organic internal standard solution was added (Ceramide/Sphingoid Internal Standard Mixture II diluted 1:10 in ethanol, Avanti Polar Lipids). The mixture was vortexed for 20 seconds and a two-phase liquid-liquid extraction was performed. The upper phase was transferred to a new clean tube and the lower aqueous phase was re-extracted with additional 2.0 ml of organic extraction solvent. The organic phases were combined and dried under nitrogen stream at 40 °C. In the case of tissue samples, the dried residue was reconstituted in 200 µl of methanol. The aqueous phase was dried in a SpeedVap solvent evaporation system and the dried residue was reconstituted in 500 µl RIPA buffer containing 5% TritonTM X-100 and total soluble protein content was determined by the BCA assay. 5 µl of reconstituted samples was injected into an LC-MS/MS system for the analysis of ceramides and sphingoid bases, 1 µl injection was required for the analysis of sphingomyelins. The system consisted of a Shimadzu LCMS-8050 triple quadrupole mass spectrometer with the dual ion source operating in electrospray positive ionization mode in the case of sphingoid bases and ceramides and in negative mode in the case of sphingomyelins. The mass spectrometer was coupled to a Shimadzu Nexera X2 UHPLC system equipped with three solvent delivery modules LC-30AD, three degassing units DGU-20A5R, an auto-sampler SIL-30ACMP and a column oven CTO-20AC operating at 40 °C (Shimadzu Scientific Instruments). Analysis of sphingolipid species was achieved using selective reaction monitoring scan mode. Sphingoid Lipid separation was achieved by reverse phase LC on a 2.1 (i.d.) x 150 mm Ascentis Express C8, 2.7 micron (Supelco) column under gradient elution, using three different mobile phases: eluent A consisting of methanol/water/formic acid, 600/400/0.8, v/v/v with 5 mM ammonium formate, eluent B consisting of methanol/formic acid, 1,000/0.8, v/v with 5 mM ammonium formate, and eluent C consisting of CH3OH/CH2Cl2 350/650. The relative concentration of each metabolite was determined using the peak-area ratio of analyte vs. corresponding internal standard. Data are reported as Analyte peak area/Internal Standard peak area and normalized according to protein content.

### C6-NBD Ceramidase Assay

C6 ceramide-NBD (ThermoFisher Scientific, N-22561) was prepared according to manufactures instruction. 7×10^4^ SUM149 cells were seeded out on coverslips in 12-well plates. Two days later, cells were washed twice in Hanks’ Balanced Salt Solution (Sigma, H8264) supplemented with 10 mM HEPES (HBSS/HEPES), before adding 5 μM C6-NBD ceramide dissolved in HBSS/HEPES. The cells were then incubated at 4°C for 30 min, washed five times in ice cold HBSS/HEPES before incubated at 37°C for 30 min. Finally, cells were washed four times before fixation with 4% PFA for 15 min. For immunofluorescence staining of the Golgi, cells were permeabilized with 0.03 µg/ml digitonin for 10 min at 4°C, and stained with RCAS1 as described above. Images were acquired using the Leica SP5 confocal microscope. ImageJ Fiji was used to measure the C6 ceramide-NBD signal in RCAS1 positive areas.

### Western Blotting

Cells for western blotting were washed once with PBS before the addition of lysis buffer (Pierce RIPA buffer, ThermoScientific, 89901). Cells were scraped in the lysis buffer supplemented with EDTA-free protease inhibitor cocktail (Sigma, 5892791001) and incubated on ice for 20 min. Lysates were centrifuged at 20800g for 15 minutes at 4°C to remove cellular debris, and protein concentration determined on the cleared lysate using the Pierce BCA Protein Assay kit (ThermoFisher, 23225). 45 μg total protein was separated by electrophoresis on NuPAGE 10% Bis-Tris protein gels (ThermoFisher Scientific, NP0315BOX) at 160 V for 1 hour and 15 min. Proteins were transferred to a PDVF membrane at 80 V for 2 hours at 4°C. Next, non-specific binding was blocked using 5% dry-milk in PBS-Tween20 (0.1%). For determining Flag-tagged PAQR4 expression, membranes were incubated with primary anti-Flag antibody conjugated to HRP for 1 hour at RT. Beta-actin served as a loading control, and membranes were incubated with primary antibody overnight. Following 3 x 5 min PBS-Tween washes, membranes were incubated with secondary antibody for 1 hour at RT. For development of the blots, washed membranes were incubated with ECL Western Blotting Substrate (ThermoFisher, 32106) and exposed to ChemiDoc XRS+ (Bio-Rad).

### Quantification and Statistical Analysis

Statistical computation on the transcriptomics data and principle component analysis on clinical ceramidase assay data were performed in RStudio. Several R packages were used to prepare the data and analyze the RNAseq. The in-house script is available upon request. For the ceramidase assays (Figure 2B, C) on the clinical patient samples, the results were analyzed by Wilcoxon matched-pairs signed rank test using GraphPad Prism 7 software. For all the comparisons, student two-tailed t test was used in GraphPad Prism 7 software. The significant symbols used in the figures are as \**P*< 0.05, \*\**P*<0.01, \*\*\**P*<0.001

### Data and Code Availability

All data and code resources supporting the current study have not been deposited in a public repository, but are available from the corresponding author on request.

### Additional Resources

There is no additional resource enclosed with the presented data.

