## Supplemental Figures for "Golgi-localized PAQR4 mediates Anti-apoptotic Ceramidase Activity in Breast Cancer"

### Supplementary Figure 1:

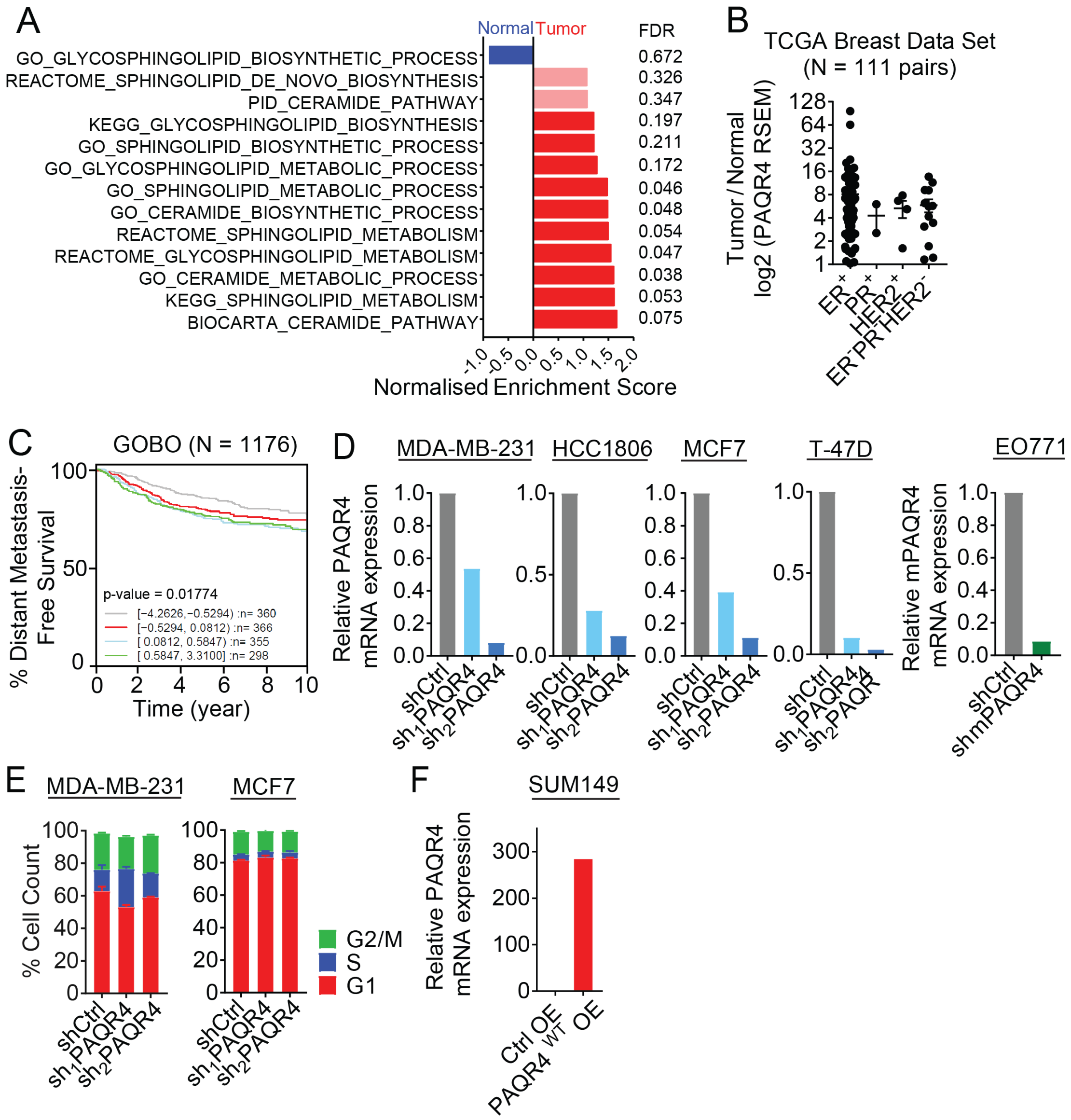


**Supplementary Figure 1: The Sphingolipid Metabolism-Related Gene *PAQR4* is Required for Breast Cancer Cellular Growth and Negatively Correlated with Breast Cancer Patient Survival.**

**A)** Gene Set Enrichment Analysis (GSEA) of sphingolipid related pathways comparing the transcriptomics of paired normal and tumor tissues from breast cancer patients (N=111) obtained from The Cancer Genome Atlas (TCGA). NES = Normalized Enrichment Score. FDR = False Discovery Rate.

**B)** Relative *PAQR4* expression in tumor compared to normal tissue samples with different estrogen, progesterone and HER2 receptor status subtypes. PAQR4 RSEM represents the normalized number of reads mapped to *PAQR4* gene.

**C)** *PAQR4* expression negatively correlates with patient survival as determined by Kaplan-Meier curves of distant metastasis-free survival based on *PAQR4* expression using Gene expression-based Outcome for Breast cancer Online (GOBO) (n = 1176 breast cancer patients) (Ringner et al., 2011).

**D)** Expression level of *PAQR4* determined by qPCR in cell lines stably transfected with shRNAs targeting *PAQR4*. *PAQR4* expression is calculated relative to control cells. The data shown is one representative of more than 3 experiments.

**E)** Cell cycle distribution analysis of MDA-MB-231 and MCF7 cells depleted for PAQR4 as determined by PI staining and flow cytometry on fixed cells. Data are presented as mean ±SEM from one of three independent experiments.

**F)** Expression level of *PAQR4* determined by qPCR in SUM149 cell lines stably transfected with retroviral overexpression system. *PAQR4* expression is calculated relative to control cells. The data shown is one representative of more than 3 experiments.

### Supplementary Figure 2:

A

```
Q86V24|PAQR2_Hs MNEPTENRLGCSRTPEPDIRLRKGHQLDGTRRGDNDSHQGDLEPILEASVLSSHHKKSSE 60
Q8N4S7|PAQR4_Hs ------------------------------------------------------------ 0


Q86V24|PAQR2_Hs EHEYSDEAPQEDEGFMGMSPLLQAHHAMEKMEEFVCKVWEGRWRVIPHDVLPDWLKDNDF 120
Q8N4S7|PAQR4_Hs -------------------------------MAFL-----AGPRLLDWASSPPHLQFNKF 24
                                 *:     .  *::     *  *: *.*

Q86V24|PAQR2_Hs LLHGHRPPMPSFRACFKSIFRIHTETGNIWTHLLGCVFFLCLGIFYMFRPNISFVAPLQE 180
Q8N4S7|PAQR4_Hs VLTGYRPA-SSGSGCLRSLFYLHNELGNIYTHGLALLGFLVLVPMTMPWGQLG-----KD 78
                :* *:**   *  .*::*:* :*.* ***:** *. : ** *  : *   ::.     ::

Q86V24|PAQR2_Hs KVVFGLFFLGAILCL------SFSWLFHTVYCHSEGVSRLF--SKLDYSGIALLIMGSFV 232
Q8N4S7|PAQR4_Hs ------GWLGGTHCVACLAPPAGSVLYHLFMCHQGGSAVYARLLALDMCGVCLVNTLGAL 132
                       :**.  *:      : * *:* . **. * :       ** .*:.*:   . :

Q86V24|PAQR2_Hs PWLYYSFYCNPQPCFIYLIVICVLGIAA-IIVSQWDMFATPQYRGVRAGVFLGLGLSGII 291
Q8N4S7|PAQR4_Hs PIIHCTLACRPWLRP-----AALVGYTVLSGVAGWRALTAPSTSARL----RAFGWQAAA 183
                * :: :: *.*          .::* :.   *: *  :::*.  .       .:* ..

Q86V24|PAQR2_Hs PTLHYVISEGFLKAATIGQIGWLMLMASLYITGAALYAARIPERFFPGKCDIWFHSHQLF 351
Q8N4S7|PAQR4_Hs RLLVFGARGVGLGSGAPGSLPCYLRMDALALLGGLVNVARLPERWGPGRFDYWGNSHQIM 243
                  * :      * :.: *.:   : * :* : *. : .**:***: **: * * :***::

Q86V24|PAQR2_Hs HIFVVAGAFVHFHGVSNLQEFRFMIGGGCSEEDAL 386
Q8N4S7|PAQR4_Hs HLLSVGSILQLHAGVV--PDLLWAAHHACPRD--- 273
                *:: *.. :  . **    :: :   .*  .
```

B

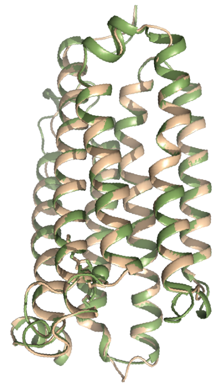


**Supplementary Figure 2: Homology Modelling Suggests that PAQR4 Function as a Ceramidase**

**A)** PAQR2 and PAQR4 amino acid sequence alignment using Clustal omega (Sievers et al., 2011). Identical residues are shown by asterisk (*). Residues with strong and weak similarities are marked with colon (:) and period (.) characters, respectively.

**B)** Overlap of predicted PAQR4 structure with the resolved crystallized structure of PAQR2.

**Supplementary Figure 3:**

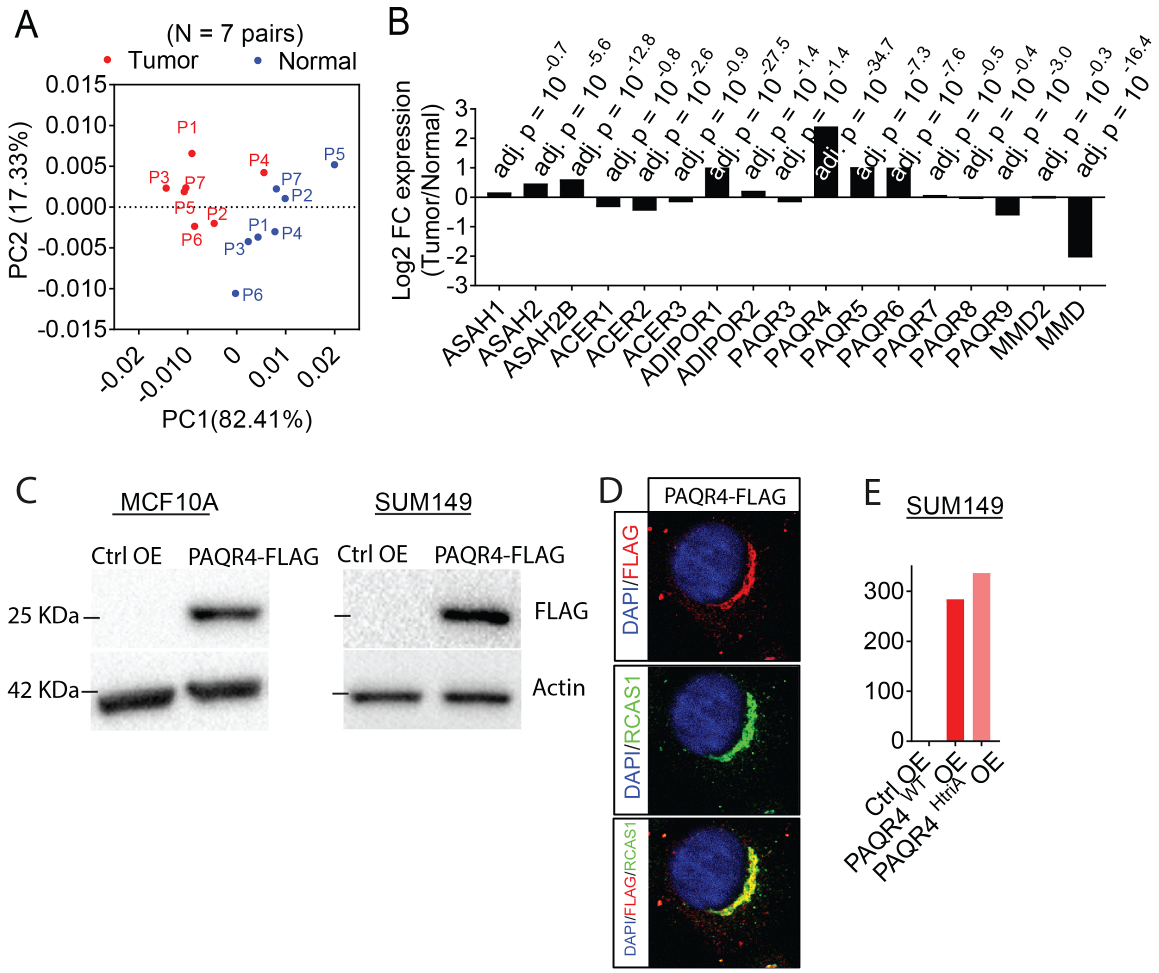


**Supplementary Figure 3: PAQR4 is a Golgi-associated ceramidase**

**A)** Principle component analysis (PCA) of the ceramidase assay on normal and breast cancer tissues. Tumor and normal tissue lysate of the breast cancer patients were incubated with deuterated-labelled ceramide species (C16, C18, C24:0 and C24:1) and the metabolites were measured by LC-MS/MS after 3 hours of incubation. The PCA on the deutrated-labelled ceramides (C16, C18, C24:0 and C24:1) and sphingosine display the clustering of tumor (red) from normal (blue) tissues.

**B)** Relative gene expression of known ceramidases in the paired breast tumor and normal tissues. The transcriptomics data were obtained from TCGA database. The adjusted p-value of the fold change in each gene expression is presented on top of each bar in the graph.

**C)** Western blot analysis of PAQR4 expression in lysates from MCF10A and SUM149 cells using anti-flag and anti-actin antibody as a loading control.

**D)** Confocal immunofluorescent images of breast cancer cell line SUM149 expressing Flag-tagged PAQR4. The tagged protein co-localizes with the Golgi marker RCAS1.

**E)** Expression level of wild type and mutated (HtriA) *PAQR4* in SUM149 cells with retroviral mediated overexpression as determined by RT-qPCR. PAQR4 expression is calculated relative to the control cells. The presented figure is one representative of more than three experiments.

**Supplementary Figure 4:**

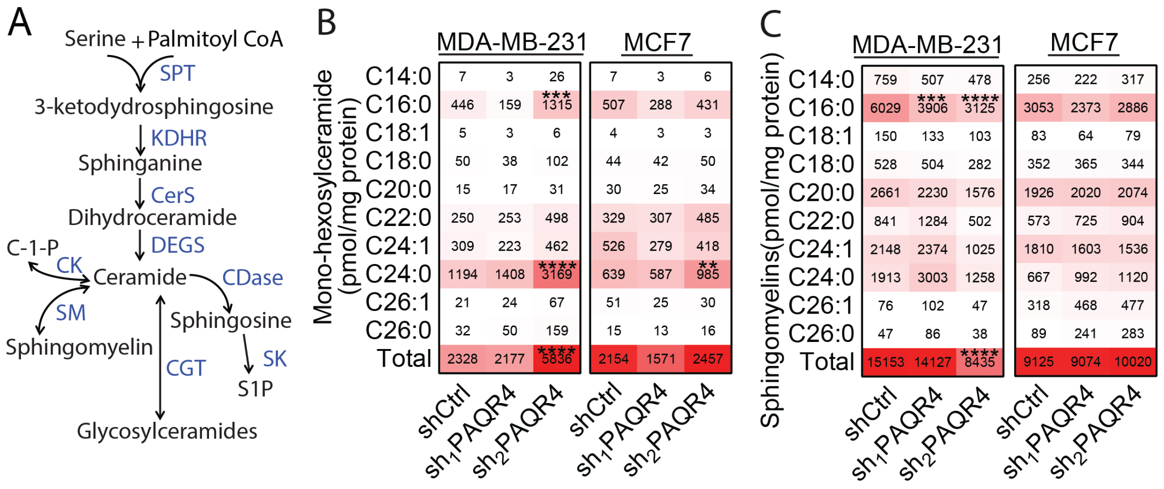

**Supplementary Figure 4: PAQR4 Depletion Causes Accumulation of *de novo* Sphingolipid Intermediates and Ceramide-induced Apoptosis**

**A)** Simplified overview of the *de Novo* sphingolipid pathway, adapted from (Mullen et al., 2012).

**B, C)** Mono-hexosylceramides and sphingomyelins levels in the lysates from MCF7 and MDA-MB-231 cells depleted for PAQR4 determined by LC-MS/MS. Sphingolipid levels were normalized to the mg protein concentration and are presented as mean values in the heatmap, n=3 per group. One out of two independent experiments is shown.

**$P$<0.01, ***$P$<0.001

**Movie S1: Molecular Dynamics Simulation of the PAQR4 Model Containing Docked a C18-Ceramide Molecule Embedded in the Phospholipid Bilayer.**
